# Mouse exploratory behaviour in the open field with and without NAT-1 EEG device: Effects of MK801 and scopolamine

**DOI:** 10.1101/2024.05.14.594089

**Authors:** Charmaine J.M. Lim, Jack Bray, Sanna K. Janhunen, Bettina Platt, Gernot Riedel

## Abstract

One aspect of reproducibility in preclinical research that is frequently overlooked is the physical condition in which physiological, pharmacological or behavioural recordings are conducted. In this study, the physical conditions of mice were altered through the attachments of wireless electrophysiological recording devices (Neural Activity Tracker-1, NAT-1). NAT-1 devices are miniaturised multichannel devices with on-board memory for direct high-resolution recording of brain activity for >48 hrs. Such devices may limit the mobility of animals and affect their behavioural performance due to the added weight (total weight of approximately 3.4 g). Mice were additionally treated with saline (control), N-methyl-D-aspartate (NMDA) receptor antagonist MK801 (0.85 mg/kg), or the muscarinic acetylcholine receptor blocker scopolamine (0.65 mg/kg) to allow exploration of the effect of NAT-1 attachments in pharmacologically treated mice. We found only minimal differences in behavioural outcomes with NAT-1 attachments in standard parameters of locomotor activity widely reported for the open field test between drug-treatments. Hypoactivity was globally observed as a consistent outcome in MK801-treated subjects and hyperactivity in scopolamine groups regardless of NAT-1 attachments. These data collectively confirm the reproducibility for combined behavioural, pharmacological and physiological endpoints even in the presence of lightweight wireless data loggers. The NAT-1 therefore constitutes a pertinent tool for investigating brain activity in e.g. drug discovery, models of neuropsychiatric and/or neurodegenerative diseases with minimal effects on pharmacological and behavioural outcomes.

## 1. Introduction

With advancements made in preclinical research, laboratories often align the mechanistic and functional aspects of animal behavioural outcomes to that of dynamic neuronal events. This is often performed with tethered or, more recently, wireless miniature recording systems [1–8] to allow recording of electroencephalography (EEG) in freely moving animals with minimal discomfort. In our laboratory, this includes the bilateral implantation of gold electrodes into the skull and attachment of a lightweight Neural Activity Tracker (NAT-1; Cybula, UK) device to this head-stage prior to behavioural testing. These devices have been employed for sleep-staging and global activity analysis during home cage recordings [7,8]. However, the surgical procedure and additional weight a mouse must tolerate amounts to about 3 g (multifunctional NAT-1 device including batteries), in addition to a small head stage modelled from dental cement and screw recording electrodes. The impact on the animal’s head could therefore conceivably limit locomotor skills and mobility of the animal and affect its natural explorative behaviour, and consequently, the outcomes of behavioural findings, thereby altering the apparent effects of pharmacological agents.

Compounds commonly used in preclinical *in vivo* research such as N-methyl-D-aspartate receptor (NMDAR) antagonist (dizocilpine; [+]-MK801) and muscarinic antagonist, scopolamine, were inoculated in mice in this study. Both compounds induce schizophrenia- [9–12] or Alzheimer-related [13,14] cognitive decline respectively and are known to induce well measurable and robust behavioural outcomes in mice including dose-dependent hyperactivity/hypoactivity [12–18]. Inclusion of pre-specified doses of MK801 and scopolamine (unpublished preliminary data) in this study will allow opposing non-systematic movements in mice to accurately represent the effects of wireless recording devices attachments in animals.

This exploratory study aims to: (1) evaluate potential differences in behavioural outcomes in the open field test in mice with and without surgically implanted head-stages/EEG recording devices (NAT-1); and (2) explore the behavioural effects of such attachments in pharmacologically treated mice. All exploratory behaviour obtainable from video-observed tests are presented to avoid bias; EEG recordings obtained in this study were not deemed relevant to changes in behavioural outcomes in mice and are therefore omitted. Generally, analyses of activity in the open field test confirm that despite very similar performance with or without EEG devices in vehicle treated control mice, subtle differences appeared in the drug groups that may inform of some behavioural alterations that affect outcomes other than activity.

## 2. Methods

### 2.1 Experimental design

The study design, analysis and reporting methods in this study were conducted in line with recommendations stated in the ARRIVE 2.0 guidelines [19] and are detailed in the relevant segments below. The study design comprised of two independent experiments conducted on separate occasions. The testing days of the week, number of animals tested per day, and testing seasons were, however, similar between studies. To ensure that any differences between Experiments 1 (no NAT-1) and 2 (with NAT-1) were the result of the attachment of wireless data logging devices but not that of variations in procedural protocols or testing environment, special care was taken to ensure that the laboratory conditions, husbandry, avoidance of bias and study designs were maintained between both experiments (detailed in Table 1). Variables which differed between Experiments 1 and 2 were marked with asterisks in Table 1 and mainly pertained to the surgical procedures and NAT-1 related handling requirements performed in Experiment 2.

**Table 1.** Details pertaining to the study designs, experimental conditions, and husbandry in Experiments 1 and 2.

|  | Experiment 1 | Experiment 2 |
| --- | --- | --- |
| **Study Outcome** | **Pharmacological studies without EEG attachments** | **Pharmacological studies in animals with EEG attachments** |
| **Persons involved (Blinder and experimenter)** | 2 | 3 |
| **Location at which tests were conducted** | Sound-attenuated room | Sound-attenuated room |
| **Animals** |  |  |
| **Strain** | C57BL/6J (Wildtype; total n = 29) | C57BL/6J (Wildtype; total n = 27) |
| **Supplier/ Breeder** | Charles River Laboratories, Kent, UK | Charles River Laboratories, Kent, UK |
| **Transport** | By road | By road |
| **Specific pathogen free conditions** | Yes | Yes |
| **Age at arrival and acclimatisation to facility** | 10 weeks; 2 weeks of acclimatisation | 9 weeks; 2 weeks of acclimatisation |
| **Housing/ husbandry; test room conditions** |  |  |
| **Cage type** | Conventional shoe box cages with wire top feeders; single housing | Conventional shoe box cages with wire top feeders (only 1 was moved to IVC cage); single housing |
| **Enrichment; bedding** | Cardboard rolls in all cages; corncob and aspen shavings (ratio 3:1) | No enrichment; corncob and aspen shavings (ratio 3:1) |
| **Temperature, humidity** | 21 ± 2 °C; 50 ± 5 % | 21 ± 2 °C; 50 ± 5 % |
| **Exposure of animal to handling** | Two/three times a day by facility staff | Two/three times a day by facility staff |
| **Noise/background noise** | Background noise (music) during light phase | Background noise (music) during light phase |
| **Dark/light cycle** | 12:12 hours; start of light phase at 07:00, no red light during dark phase | 12:12 hours; start of light phase at 07:00, no red light during dark phase |
| **Diet** | Normal chow pellets (SDS CRM) and room-temperature water *ad libitum* | Normal chow pellets (SDS CRM) and room-temperature water *ad libitum* |
| **Handling of cages** | Cleaning once every 2 weeks; water and food changed/topped up twice a week. | Cleaning once every 2 weeks; water and food changed/topped up twice a week. |
| **Experimental: animals** |  |  |
| **Age of animal at start to experiment** | 12-13 weeks (at the start of testing) | 10-11 weeks (at surgery) and 12-13 weeks (at the start of testing) |
| **Transport of animals in facility** | Silent trolley type; cages were not blacked out or covered during transport | Silent trolley type; cages were not blacked out or covered during transport |
| **Time of testing, day of week** | Commencement at 10:00; testing was conducted from Monday to Thursday on the first week, and Monday to Wednesday on the second week. | Commencement at 10:00; testing was conducted from Monday to Thursday on the first week, and Monday to Wednesday on the second week. |
| **Total duration of test phase per day** | Approx. 5 hours (4 animals tested am, 3 tested pm) | Approx. 5 hours (3 animals tested am, 3 tested pm) |
| **Animal under diet restrictions during experimental phase** | No | No, except during NAT-1 application and drug administration: water bottles were removed, and cage lids were overturned to prevent NAT devices from accidentally wedging onto cage-lids and food-hoppers |
| **\*Naivety to experimental paradigm and drugs** | Naïve to both experimental procedures and drugs | Home cage EEG recording and check, 24 hours prior to open field test; naïve to drugs |
| **Experimental: surgical protocols** |  |  |
| **\*Time period between surgeries and start of experiment** | N.A. | 2 weeks |
| **\*Total duration of surgery per day** | N.A. | Approx. 7 hours/day (commencement at 08.00; surgeries performed only on Mondays and Tuesdays |
| **\*Anaesthetic** | N.A. | Gas anaesthesia (isoflurane) 1.0 ml saline (intraperitoneal) and 0.2 ml buprenorphine (subcutaneous; Vetergesic®, Cera Animal Health Ltd, UK). |
| **\*Analgesia** | N.A. | Drinking water was supplemented with Rimadyl® for 2 days before and after the surgery |
| **\*Surgical procedure** | N.A. | 1 surgeon; 2 assistants |
| **Experimental: design and methods/data analysis** |  |  |
| **Drug, and concentration** | 0.9 % saline; 0.65 mg/kg MK801; 0.85 mg/kg scopolamine. Intra-peritoneal administration of 10mg/kg of treatment. | 0.9 % saline; 0.65 mg/kg MK801; 0.85 mg/kg scopolamine. Intra-peritoneal administration of 10mg/kg of treatment. |
| **\*Experimenter** | 1 female | 2 females and 1 male |
| **\*Frequency of handling** | Handling during drug injections, immediately before and after test. | Handling during NAT-1 applications, drug injections, immediately before and after test. |
| **Randomisation method for allocation of animals to test groups** | random.org, Williams square design. | random.org, Williams square design. |
| **Blinding of experimenter to group allocation** | Yes. | Yes. |
| **Standardisation of animal handling across experimenters (protocol and training)** | Yes (1 in randomisation and blinding, 1 in drug administration and experiment) | Yes (2 involved in NAT-1 device application and drug administration and randomisation, 1 in experiment) |
| **Software used in recording and data extraction** | Automatic tracking software, ANY-Maze (v.5.1) | Automatic tracking software, ANY-Maze (v.5.1) |
| **Inclusion and exclusion criteria** | No inclusion criteria. Exclusion criteria based on analysis using Grubb's method before re-evaluation of animal behaviour (video). | No inclusion criteria. Exclusion criteria based on analysis using Grubb's method before re-evaluation of animal behaviour (video). |
| **Statistical analysis** | Bootstrapped one-way ANOVA and t-test for conventional analysis; repeated measures ANOVA for binned analysis; and estimation statistics to determine the magnitude of differences. | Bootstrapped one-way ANOVA and t-test for conventional analysis; repeated measures ANOVA for binned analysis; and estimation statistics to determine the magnitude of differences. |
| **Experimental: complexity of test arena** |  |  |
| **Test arena** | Square; 50 x 50 x 50 cm; white Perspex | Square; 50 x 50 x 50 cm; white Perspex |
| **Average light intensity in test arena** | 320 lux | 320 lux |
| **Temperature and humidity in test room** | 20 ± 2 °C; 48 ± 5 %; no background noise | 20 ± 2°C; 48 ± 5 %; no background noise |
| **Number of test arenas used at any given time** | 1 | 1 |
| **Cleaning agents** | Non-fragrant and alcohol-free wet wipes | Non-fragrant and alcohol-free wet wipes |
| *Inconsistent experimental factors which differed between experiments. |  |  |

### 2.2 Animals

Twenty-nine C57BL/6J (10-11 weeks old male mice) from Charles River Laboratories (Kent, England) were obtained for Experiment 1 and 27 for Experiment 2. Mice were housed individually in standard Macrolon shoe box cages (Techniplast 1145) measuring at 369 x 165 x 132 mm, under a 12- hour light/12-hour dark cycle (lights on at 7am). Temperature and humidity were set at 20 ± 5 °C and 48 ± 5 % in both housing and experimental rooms. Animals had free access to standard rodent food chow (SDS 801190 RM3, England) and water *ad-libitum*. All animals were acclimatized to the facility environment and extensively handled for 2 weeks prior to the start of surgeries and/or open field testing. Mice were tail-handled to avoid detachment of the head-stages and management of severe stereotypic responses during transfer. All housing and handling of animals were in accordance with the international standards on animal welfare (EU/63/2010) and Home Office (UK) regulations.

### 2.3 Stereotactic surgery and home-cage recordings (Experiment 2 only)

Animals had free access to water supplemented with carprofen Rimadyl^®^ (5mg/kg; Zoetis, UK) and were acclimatised to soft diet (Transbreed (E), SDS 801222, England; and Solid Dirink® SDST-75, Netherlands) 48 hours prior to and after surgery. Stereotaxic surgeries were conducted as described previously (Crouch et al., 2018, 2019; Jyoti et al., 2010; Platt et al., 2011). Briefly, anaesthesia was induced with 3 % isoflurane (IsoFlo^®^, Zoetis) in medical grade oxygen and maintained on 1.5 % isoflurane during surgery. Mice were held in a stereotaxic frame (Stoelting, Wood Dale, Il, USA). Epidural gold-plated screw electrodes were placed at the following stereotaxic locations [20] on exposed skulls from the prefrontal cortex (2 mm anterior to bregma/close to midline), left and right hippocampus (2 mm posterior to bregma, 1.5 mm lateral to midline). Reference and ground electrodes were placed in a neutral location above the occipital plate. Electrodes were soldered and assembled into a 6-pin adaptor and fixed on to the skull by a mixture of Durelon^®^ dental cement and tissue adhesive. The pins measured 0.40 mm (square) with spacing pitch of 1.27 mm. Once the dental cement dried, the animal was removed from the stereotaxic instrument and injected with 1.0 ml saline (intraperitoneal) and 0.2 ml buprenorphine (subcutaneous; Vetergesic^®^, Cera Animal Health Ltd, UK). Animals were placed on heating pads immediately after the procedure and further analgesic treatment continued for 2-3 days as appropriate. Following surgery, animals were weighed daily to monitor their recovery and were provided with a soft diet. A minimum of 7 days was allowed for recovery before the start of testing. Nine animals did not recover 48 hours post-surgery or had dislodged head-stages and were euthanised.

### 2.4 Drugs and drug formulation

(+)-MK801 maleate (CAS: 77086-22-7) and scopolamine hydrobromide (CAS: 114-49-8) were obtained from Tocris Bioscience (Bristol, UK) and diluted to 0.65 mg/kg MK801 and 0.85 mg/kg scopolamine (salt-corrected for both drugs) in 0.9% saline 24 hours prior to intraperitoneal injections. Saline was administered as vehicle-control. Reagents were administered (10 ml/kg) as a single dose, with a sterile injection needle, 30 minutes prior to behavourial testing. Assignment of all mice to different drugs were randomized (Experiment 1: n = 10 for saline, n = 9 for MK801, and n = 10 for scopolamine; Experiment 2: n = 9 per drug group) using a Williams Square design [21].

### 2.5 Apparatus and testing protocols

Upon behavioural testing, mice were aged 12-13 weeks old and weighed approximately 26.8 ± 2.0 g in the open field arena. Intraperitoneal administration of drugs and testing sequence was conducted in a blinded manner. In Experiment 2, mice were drug-treated immediately before the attachment of NAT-1 devices. NAT-1 measures 18 x 22.2 x 10.3 mm and weighs approximately 2.2 g including a zinc battery (Zinc-air, size 13, Duracell EasyTab). The total weight of the NAT-1 and head- stage (ranging between 1.1 and 1.5 g) equals approximately 3.4 g. Common to both experiments, subjects were habituated to the test room for 30 minutes, following drug inoculations.

To quantify the behavioural outcomes of using the NAT-1, we tested activity of all mice in the open field test [22–24]. The open field tests were conducted in a dedicated sound-attenuated room and in a square box (made up of white reflective Perspex) of 50 x 50 x 40 cm (length x width x height). Two white LED lights facing upwards from the arena and the main fluorescent room lights were switched on, giving rise to an average illumination of 320 lux. Temperature and humidity were maintained at 20 ± 5°C and 48 ± 5% respectively. An overhead camera (Imaging Source, DMK22AUCO3), 125 cm from the floor of the arena, was connected to an automated recording system (ANY-Maze v. 5.1, Stoelting) for online tracking of the animal’s centre of mass. Each mouse was placed into the centre of the open field and allowed to freely explore for 30 minutes, and the apparatus was thoroughly cleaned with odourless wet wipes between each mouse. The automatic tracking option was defaulted in ANY-Maze to allow the software to adjust tracking parameters by contrasting the apparatus background, illumination, and animal coat colour. Sampling rates were maintained at the programme’s recommended 15 frames per second (fps) for small animals. All analyses were conducted offline with the same software (ANY-Maze v. 6.1). Parameters which were widely recorded by different laboratories (see [25] for review) were explored. These included: *total distance travelled in the arena*, *line-crossings in a 16-grid overlay*; *meandering* (a measurement of an animal’s *absolute turn angle* over the *total distance travelled* which refers to the winding movement of animal); *frequency of rotations* (total of 360 degree turns to the left and 360 degree turns to the right); *degree of thigmotaxis* (a ratio of *distance travelled in the periphery zone* against the *total distance travelled*); and the *average distance from the centre-point* (to provide meaningful indications of the general location of the mice without utilizing zonal outlines). Thigmotaxis zone was defined as the zone 5 cm from the arena walls.

### 2.6 Exploratory data analyses

Representative track-plots of animals in each drug group of Experiments 1 and 2 were generated by extracting pixel values of the animal’s centre point from ANY-Maze and converting them to millimetres using a MATLAB script. The track-plots were further divided into two-time intervals (0- 15 minutes and 15-30 minutes) for clarity and plotted using MATLAB (R2019a, Mathworks, USA).

Additional exploratory analyses of all outcomes obtainable from ANY-Maze were conducted to minimise and avoid hypothesis-driven selection of parameters in the open field test [26] by means of heat-mapping [27,28]. A total of 68 analysis parameters were extracted by ANY-Maze and were listed for comparison. Heat-maps were created to show all data points for the comparison of effects of NAT- 1 on each drug condition. The heat-maps were generated with *p*-values obtained from independent two-tailed T-tests with Satterthwaite correction for heteroscedasticity and bootstrapped 1000 times with replacement (seed set at 123456). The intensities of colours on the heat-map were scaled according to the *p*-value threshold set at 95% confidence level. Parameters in the heat-maps were clustered for clarity. *Apparatus measures* (#1-20) concerning *total distance* and other additional information regarding activity for the entirety of the test were categorized at the top of the map. *16-grid crossings* (#21), *centre-point* measures (#22-29) and *thigmotaxic zone* measures (#30-68) which are generally used as *standard parameters* were clustered next (Table S1). Heat-maps represent global read-outs over 30 minutes so time-dependent differences and scoring by *segment of test* were not considered. *Immobility* and *freezing* parameters were defaulted in ANY-Maze as 2000 milliseconds (sensitivity level set at 65%) and 1000 milliseconds (threshold levels on at 30 and off at 40) as the minimum immobility and freeze durations respectively, and no changes were made to both criteria. Parameters from head and tail tracking were not recorded.

### 2.7 Statistical analysis

Parameters of interest obtained from the tracking software and calculated parameters of interest (e.g. total rotations, meandering and thigmotaxis) were first tested for normality using Lilliefors Kolmogorov-Smirnov test. The observed unpaired mean differences were reported and derived from the expected sampling error to illustrate the magnitude of differences using estimation analysis [29]. T-tests, or analysis of variance (ANOVA; for total responses parameters) with post-hoc Bonferroni’s corrections were performed for all conventional p-value analyses and bootstrapped 1000 times with replacement (seed set at 123456) to accommodate non-normally distributed outcomes [30]. Group differences were also considered negligible if effect sizes (ηp^2^) were less than 0.1. Only data for distance moved for the entirety of the test were further collapsed into 5-minute bins; statistical differences were evaluated using two-way ANOVA for repeated measures, with experimental groups as between-subject factors and time as within-subject factors. Statistical significance was set at 95% confidence level for conventional statistics. Outliers (with residuals more than two standard deviations away from the mean) were re-investigated for extreme experimental issues or recording errors. No convincing reasons to remove any data on this basis were found. All conventional analysis, analysis of estimates and visualisation of outcomes were performed in R (v.4.3.0; [31]).

## 3 Results

The experimental and housing conditions for both independent experiments are detailed in Table 1; the surgical procedures and NAT-1 attachments were the only inconsistent variables between both experiments. Comparisons for each drug treatment were initially visualised in a heat-map (Figure 1); significant differences where p<0.05 thresholded and are represented in blue. Head-stage/NAT-1 attachments led to significant differences in 32 parameters for the saline-treated groups, 12 parameters for the MK801-treated groups and 14 parameters for the scopolamine-treated groups. No parameter showed a statistical significance in all comparisons between experiments for all treatments (Figure 1 and Table S1).

**Figure 1.**
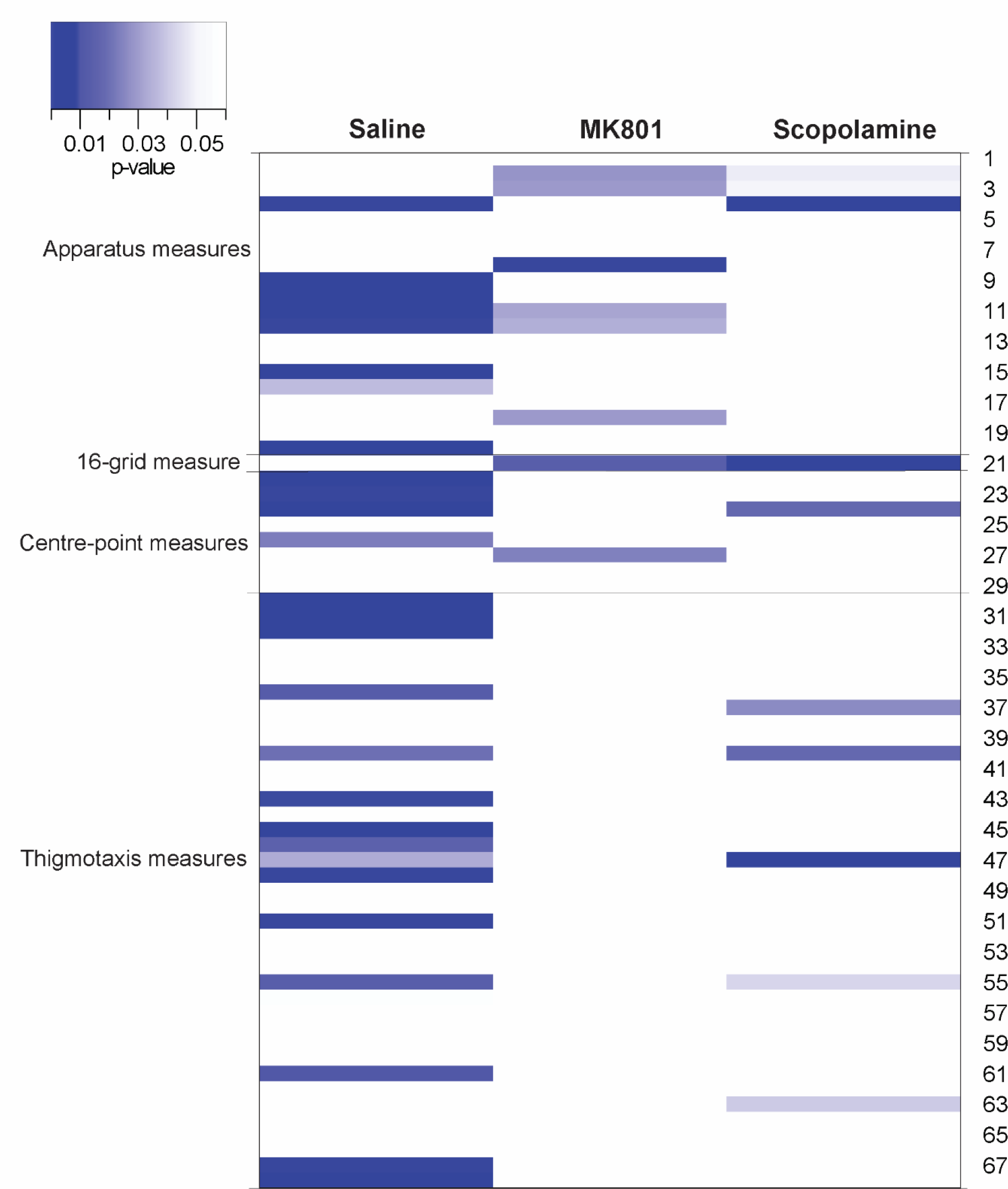
Heat-map comparing experiment 1 (no head-stage/NAT-1) and 2 (with head-stage + NAT-1) for each drug treatment. Parameters are clustered into *apparatus measures* (#1-20; commonly used in the analysis of standard parameters); *16-grid measure* (#21); *centre-point measures* (#22-29); and *thigmotaxis measures* (#30-68). The intensity of colours on the heat-map was scaled accordingly with the head- stage/treatment cohorts and the parameter presenting the greatest differences of *p*-values between head-stage/treatment cohorts were isolated and analysed. Data presented in blue in the heat-map represents a *p*-value with the threshold of 0.05 in the between-experiment comparisons. The heat-map was visualized in R (v4.3.0).

The explorative strategies were visualised by track-plots of representative animals in Experiments 1 and 2 (Figure 2a). Locomotor activity and spatial distribution in the open field were maintained between animals with or without NAT-1 of the same drug treatment in both the first and the second 15-minute interval of the experiment. For both experiments, scopolamine-treated animals remained in the *periphery* whilst the MK801-treated animals remained in a section of the arena, generally localized away from the arena walls. The saline-treated animals traversed across the centre with a greater frequency than the two drug groups and hence did not show a clear preference for an exploration zone.

**Figure 2.**
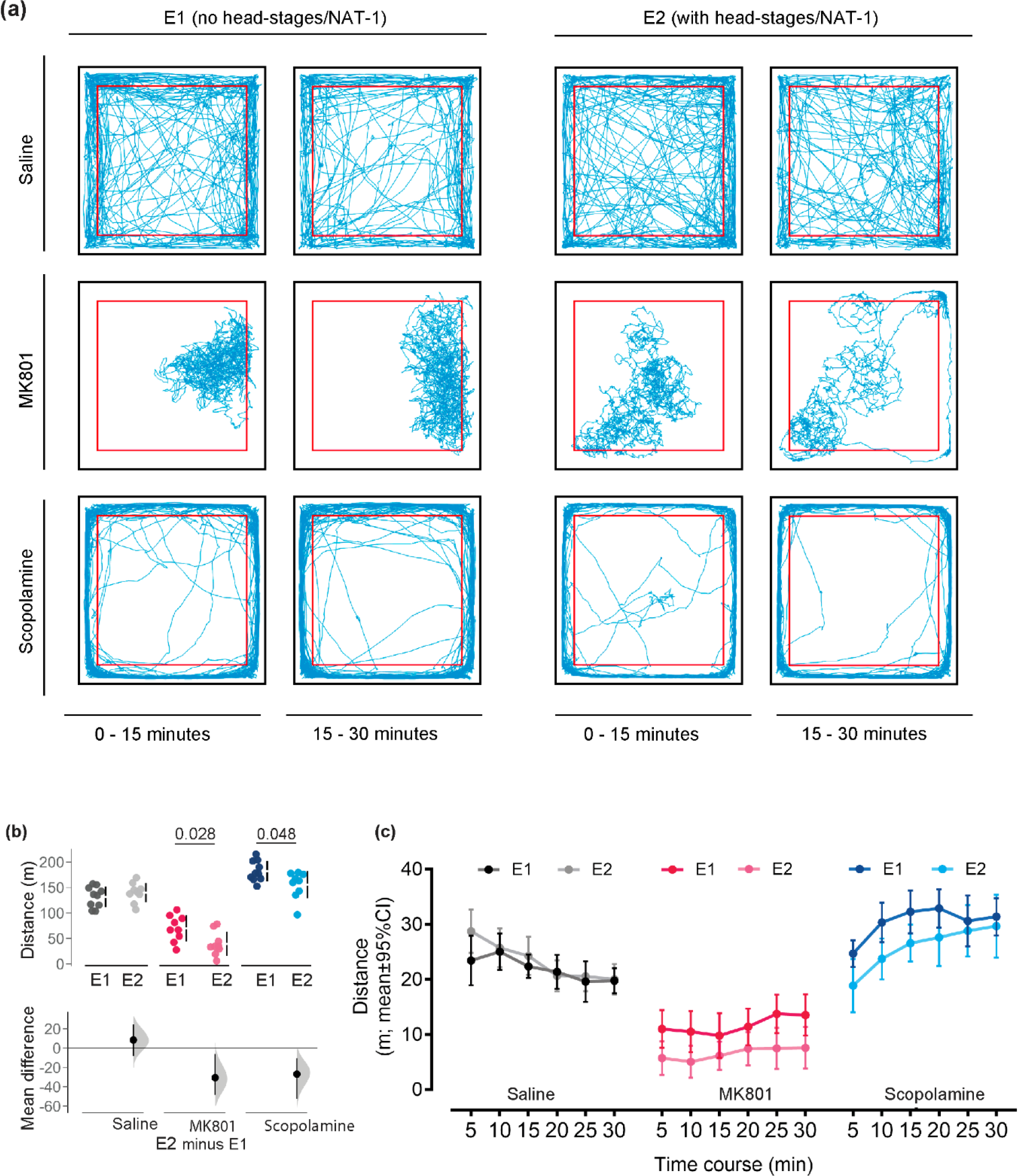
Behavioural responses for locomotor activity between Experiments 1 (E1) and 2 (E2). (a) Representative exploration paths of animals (with and without head-stage + NAT-1) administered with vehicle, MK801 or scopolamine recorded in the open field test and visualized by track-plots divided into time bins, 0-15 minutes and 15-30 minutes. The *periphery* (thigmotaxis zone) of the open field arena is demarcated by a red border, 5 cm away from the walls of the arena. Representative paths were illustrated using an in-house MATLAB (R2018a, Mathworks, USA) script. (b) Analysis of locomotor activity by *cumulative distance travelled* and (c) *total distance travelled* over 30 minutes of testing in the open field was visualised in 5-minute time bins. Individuals are represented by single data points with their respective means (gap) ± standard deviation (vertical lines) in the top axis. The shaded curve and the error in the bottom axis demonstrate the distribution of sampling error and its respective 95% confidence interval for the difference between the means. Performance of animals in E1 is represented by the 0 line in the bottom axis. Conventional statistics (independent sample t-tests) revealing a significance set at 95% (*p*<0.05) are annotated as text on (b). All analyses were bootstrapped at 1000 resamples, with seed set at 123456 and visualized in R (v.4.3.0). E1, with no head-stage/NAT-1; and E2with head-stage and NAT-1.

The head-stage/NAT-1 attachments decreased *total distance travelled* by 30.6 m [95% CI -48.6, -6.04 m] and 26.9 m [95% CI -52.3, -10.4 m] in MK801-and scopolamine- treated animals respectively. Means were outside of the interval estimate, but minimally increased *total distance travelled* by 8.37 m [95% CI -8.09, 24.6 m] in the saline-treated group (Figure 2b). *Total distance travelled* analysed in 5-minute time intervals are visualised in Figure 2c. No main effect of head-stages was observed for saline- treatments, but significant reductions were yielded for MK801- (F(1,16)=6.98, *p*=0.018, ηp^2^=0.304) and scopolamine- (F(1,17)=6.26, *p*=0.023, ηp^2^=0.269) treated cohorts.

Comparisons of animals with and without NAT-1 are statistically summarised in Table 2 and visualised in Figure 3. No notable differences due to NAT-1 attachments were discerned in the *average distance from the centre-point* (m [95% CI]: saline, -0.020 [-0.026, -0.012]; MK801, 0.006 [-0.041, 0.036]; and scopolamine, -0.080 [-0.018, 0.030]) ; Figure 3a). *Line-crossings* differed between experimental groups for all treatments (Figure 3b)- NAT-1 attachments induced the largest reductions in line-crossing counts in MK801- (-409 counts [95% CI, -600, -158 counts]) and scopolamine- (-362 counts [95% CI,-541, -242 counts]) treated animals, but minimally affected saline-treated mice (-28.6 counts [95% CI, -148, 86.7 counts]) .

**Figure 3.**
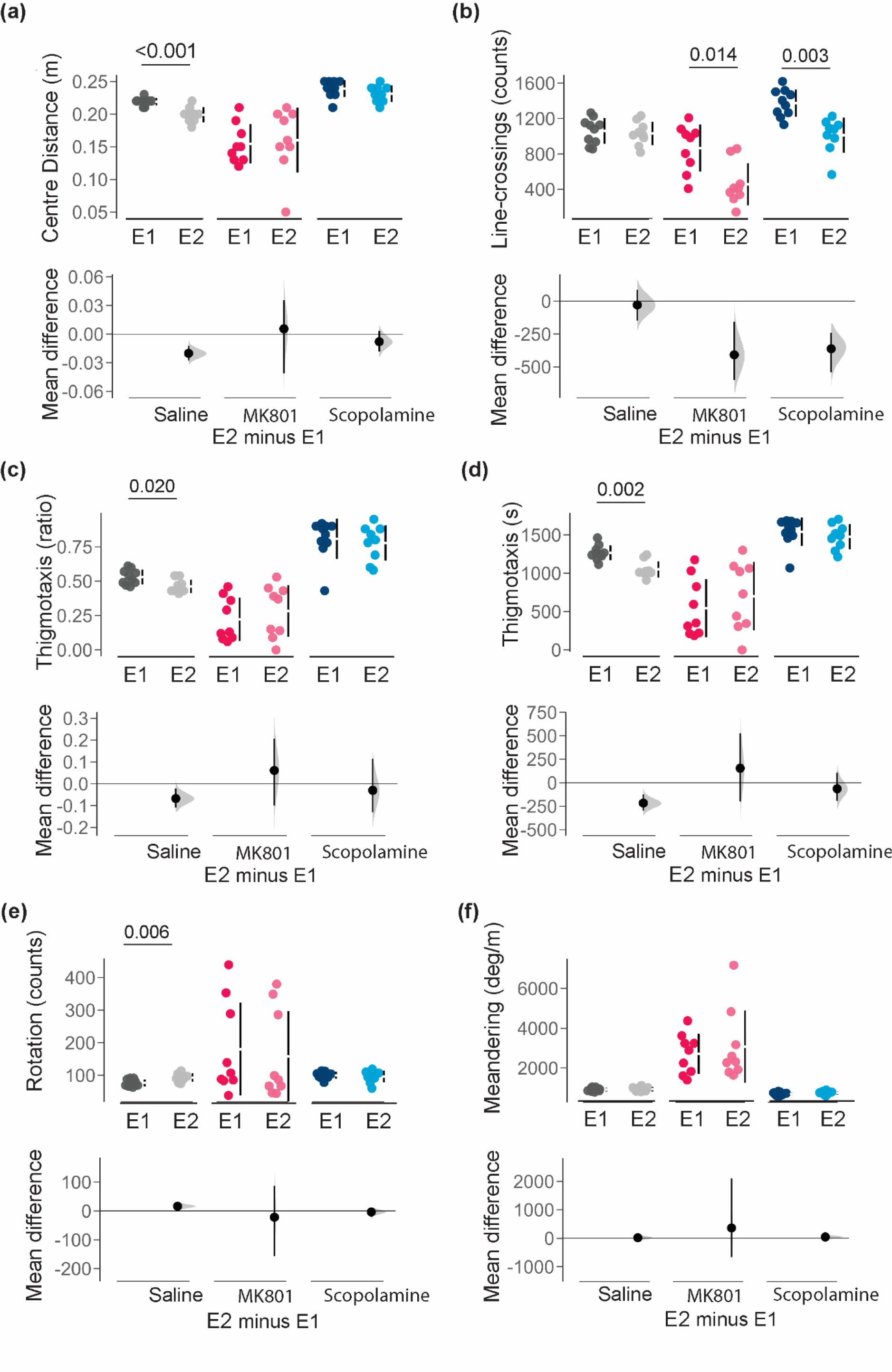
Comparison of behavioural responses for parameters of distance between Experiments 1 (E1) and 2 (E2). (a) *average distance from the centre-point*; (b) *line-crossings*; (c) *thigmotaxic ratio*; (d) *time spent in the thigmotaxic zone*; (e) *total number of rotations*; and (f) *meandering*. Individuals are represented by single data points with their respective means (gap) ± standard deviation (vertical lines) in the top axis. The shaded curve and the error in the bottom axis demonstrate the distribution of sampling error and its respective 95% confidence interval for the difference between the means. Performance of animals in E1 is represented by the 0 line in the bottom axis. Conventional statistics (independent sample t-tests) revealing significance set at 95% (*p*<0.05) are annotated as text on the graphs. All analyses were bootstrapped at 1000 resamples, with seed set at 123456, and visualized in R (v.4.3.0). E1 with no head- stage/NAT-1; and E2 with head-stage and NAT-1.

**Table 2.** Summary of between-experiment comparisons for each drug treatment in pre-selected parameters. Data was obtained by bootstrapped independent T-tests (two-tailed) and mean difference was obtained using estimation analysis. All analyses were resampled 1000 times (seed starting at 123456) and obtained in R (v.4.3.0) and significance was considered where *p*<0.05 (highlighted in red).

| Parameters | Drug | Between-experiment comparisons |  |  | E2 minus E1 |
| --- | --- | --- | --- | --- | --- |
| | | E1 | E2 | $p$-value | Mean difference<br>[95% CI] |
| | | mean±SD | mean±SD | | |
| Distance moved (m) | Saline | 131.6±20.2 | 139.9±19.2 | 0.347 | 8.4 [-8.1; 24.6] |
| | MK801 | 70.0±25.6 | 39.3±23.5 | 0.028 | -30.6 [-48.6; -6.0] |
| | Scopolamine | 182.2±20.0 | 155.2±26.7 | 0.048 | -26.9 [-52.3; -10.4] |
| Average distance from centre-point (cm) | Saline | 0.218±0.007 | 0.199±0.010 | 0.001 | -0.020 [-0.026; -0.012] |
| | MK801 | 0.155±0.030 | 0.161±0.050 | 0.782 | 0.006 [-0.041; 0.036] |
| | Scopolamine | 0.237±0.013 | 0.230±0.012 | 0.275 | -0.008 [-0.018; 0.003] |
| Thigmotaxis (ratio) | Saline | 0.532±0.052 | 0.463±0.049 | 0.020 | -0.068 [-0.11; -0.021] |
| | MK801 | 0.223±0.157 | 0.285±0.189 | 0.471 | 0.061 [-0.1; 0.207] |
| | Scopolamine | 0.807±0.147 | 0.777±0.128 | 0.667 | -0.030 [-0.131; 0.115] |
| Thigmotaxis (s) | Saline | 1266.1±96.3 | 1050.0±105.7 | 0.002 | -216 [-295; -123] |
| | MK801 | 545.1±379.3 | 699.2±445.6 | 0.435 | 154 [-199; 527] |
| | Scopolamine | 1539.8±184.5 | 1475.8±163.8 | 0.437 | -64 [-192; 107] |
| Line-crossings | Saline | 1060.6±145.3 | 1030.0±132.49 | 0.442 | -28.6 [-148; 86.7] |
| | MK801 | 866.9±265.1 | 458.3±238.5 | 0.014 | -409 [-600; -158] |
| | Scopolamine | 1374.2±154.3 | 1012.0±196.9 | 0.003 | -362 [-541; -242] |
| Meandering (degrees/s) | Saline | 912.5±89.5 | 932.0±107.4 | 0.668 | 19.5 [-63.6; 102] |
| | MK801 | 2716.4±1008.4 | 3077±1813.4 | 0.603 | 361 [-663.0; 2100] |
| | Scopolamine | 705.6±73.5 | 750.4±89.2 | 0.325 | 44.7 [-25.6; 117] |
| Rotation | Saline | 76.5±10.5 | 98.8±12.5 | 0.006 | 16.3 [5.6; 26] |
| | MK801 | 180.1±142.7 | 158.2±138.3 | 0.758 | -21.9 [-156.0; 87.5] |
| | Scopolamine | 99.8±10.6 | 96.1±18.2 | 0.623 | -3.7 [-18.9; 7.16] |

Thigmotaxis ratio was calculated by thigmotaxis distance divided by total distance moved; SD, standard deviation; CI, confidence intervals (upper, lower bounds); E1, Experiment 1 with no head-stage/NAT-1; and E2, Experiment 2 with head-stage/NAT-1.

In parameters of localization, head-stage/NAT-1 attachment in animals was estimated to decrease *thigmotaxis ratio* and *thigmotaxis time* in saline-treated animals (-0.068 [95% CI, -0.11, -0.021] and -216 s [95% CI, -295, -123 s] respectively), but not under exposure to MK801 or scopolamine (Figure 3c and d). Comparisons between Experiments 1 and 2 also confirmed minimal differences in *rotations* (counts [95% CI]: saline, 16.3 [5.6, 26]; MK801, -21.9 [-156.0, 87.5]; and scopolamine, -3.7 [-18.9, 7.16]) or *meandering* (degrees/m [95% CI]: saline, 19.5 [-63.6, 102]; MK801, 361 [-663, 2100]; and scopolamine, 44.7 [-25.6, 117]) with NAT-1 attachments in all mice (Figure 3f).

The trend of activity in MK801- and scopolamine-treated animals (relative to saline-treated mice) is robust for both experimental groups. Within-experiment analyses also showed that MK801- treated animals had decreased, global activity in the open field (relative to saline-treated animals; Table S2), independent of NAT-1 attachments. The contrary was observed for scopolamine-treated animals.

## 4. Discussion

The primary objective of this study was to evaluate the robustness of behavioural findings between animals with and without head-stage/NAT-1 attachments in the open field. This device differs in many respects from other implantable miniaturised telemetric EEG recording systems for mice [32–35] and is considered an advancement to many tethered recording devices (see Platt et al., 2011 for discussion). While many of these systems are widely used, a comparison between unoperated and surgically operated and subjects are not commonplace but important to define the robustness of behaviours under scrutiny. For the open field, the behavioural repertoire in saline-treated animals were shown to be generally maintained across experimental groups with minimal implications of the surgery, head-stage moulding and NAT-1 attachments. Animals treated with MK801 or scopolamine on the other hand responded to these modifications with an overall lowering of locomotor activity, with less wall hugging and more frequent visits to the centre of the arena. This suggests that despite considerable weight, NAT-1 per se is not inducing behavioural changes, but these can be visualised in the presence of drugs. Such behavioural anomalies may include stereotypy, but most obviously were reflected in hypoactivity in this study. Despite wider confidence intervals in the NAT-1 cohorts treated with either MK801- or scopolamine- treatment the magnitude of differences between drug groups and controls were in the same direction and numerically very similar, supporting the robustness of the data, and the resilience of the mice against neurosurgical intervention and weight bearing head-attached devices.

We therefore confirm that lightweight untethered head-stage/NAT-1 attachments do not impose serious limitations on locomotor activity- the primary outcome commonly assessed in the open field. However, there were variations pertaining to parameters of movement types and location-specific proxies, particularly in the thigmotaxic zone. Differences in location-specific proxies can potentially be explained by the animals’ somatosensory perception of the device on their head (‘object permanence’; [36,37]. The device may present a steric hinderance for the mobility of the animals as indicated by reductions in thigmotaxis measures in the wall zone of the arena. Other non-prespecified endpoints in this study showed variable NAT-1 attachment-dependent differences and included parameters of ethological values such as *freezing* or *heading error*, *the zone which was first entered*, *path efficiency to zone*, and *corrected integrated path length*. Intriguingly, drug dependent changes in these proxies were not influenced by NAT-1 attachments.

Surgical intervention in Experiment 2 is arguably the biggest differential that may contribute to irreproducibility in behavioural outcomes between groups. It is imperative to consider possible side-effects of the implanted electrodes or surgical process which may provoke changes in sleep pattern [38]; induce meningeal lymphangiogenesis and affect brain homeostasis [39]; or increased microglial activation and seizure susceptibility [40]. Weight bearing and drug treatment may further exacerbate such side-effects (see [41] for discussion). What appears crucial is that a direct comparison between unoperated and surgically operated specimens with telemetric devices should be performed for each behavioural test before subjects with EEG recording devices can be classed as ‘normal’ controls. There is still abundant room for further assessments to determine if responses remain robust in more complex behavioural paradigms or in e.g. other mouse lines or genetically modified strains. Aspect that may be addressed in the future is the effect of head-stage/NAT-1 attachments on more fine-grained behavioural fractionations using unsupervised, data-driven approaches [42,43] which require considerable hardware and software upgrades. . This may be relevant to studies exploring severe behavioural anomalies in different etiological responses as indications of anxiety [44–46] or epilepsy [47].

Overall, our data provide compelling evidence for by and large undisturbed behaviour in terms of spatial exploration, as all changes induced by drugs in reported parameters were similarly observed with and without NAT-1 attachment. This study confirms the robustness of behavioural responses in the context of using wireless recording devices in the open field. Data robustness *per se* is important as many laboratories provide EEG findings aligned with behavioural responses, such as ours [3,7,48]. Normative behaviour is critical if EEG recordings are interpreted as representative for a ‘normal’ animal.

## Patents

Not applicable.

## Acknowledgements

We would also like to acknowledge the staff of the Medical Research Facility, University of Aberdeen, for their support with animal care, handling and behavioural experiments.

## Author Contributions

**Charmaine Lim**: Conceptualization, Methodology, Investigation, Formal Analysis, Data archiving, Writing – original draft; **Jack Bray**: Methodology, Investigation; Review; **Sanna Janhunen**: Conceptualization, Methodology, Review & Editing; **Bettina Platt**: Conceptualization, Methodology, Supervision, Review and Editing; **Gernot Riedel**: Conceptualization, Project administration, Supervision, Methodology, Writing – Review & Editing, Funding. All authors have read and agreed to the published version of the manuscript.

## Institutional Review Board Statement

The animal study protocol was approved by the Institutional Review Board (or Ethics Committee) of the University of Aberdeen.

Informed Consent Statement: Not applicable.

## Data Availability Statement

All data are presented in this paper and numerically will be made available on request.

## Supplement

**Table S1:** List of obtainable parameters in ANY-Maze with their associated *p-*values from one-way analysis of variance and post-hoc analysis with Bonferroni’s corrections (between-drug comparisons in each experiment) and independent T-test analysis (between-experiment comparisons; corresponding heat-map found in Figure 1). Significant associations were *p*<0.005 are highlighted in red and significant associations common to both experiments are highlighted in red and bolded. Statistical analyses were obtained in R (v.4.3.0).

| Number and parameter name |  | Between-drug comparisons<sup>a</sup> |  |  |  | Between-experiment comparisons<sup>b</sup> |  |  |
| --- | --- | --- | --- | --- | --- | --- | --- | --- |
|  |  | Experiment 1 |  | Experiment 2 |  | Saline | MK801 | Scopolamine |
|  |  | MK801<br>(ref. saline) | Scopolamine<br>(ref. saline) | MK801<br>(ref. saline) | Scopolamine<br>(ref. saline) |  |  |  |
| Apparatus measure |  |  |  |  |  |  |  |  |
| 1 | Duration | 1.000 | 1.000 | 1.000 | 1.000 | 1.000 | 1.000 | 1.000 |
| 2 | Distance | 0.000 | 0.000 | 0.000 | 0.531 | 0.347 | 0.028 | 0.048 |
| 3 | Mean speed | 0.000 | 0.000 | 0.000 | 0.493 | 0.357 | 0.029 | 0.050 |
| 4 | Max speed | 0.000 | 1.000 | 0.000 | 1.000 | 0.001 | 0.130 | 0.004 |
| 5 | Freezing episodes | 0.064 | 0.136 | 0.042 | 1.000 | 0.079 | 1.000 | 1.000 |
| 6 | Time freezing | 0.120 | 0.263 | 0.481 | 1.000 | 0.077 | 0.553 | 0.430 |
| 7 | Freezing latency | 1.000 | 1.000 | 0.690 | 1.000 | 0.732 | 0.239 | 0.359 |
| 8 | Mean freezing score | 0.938 | 1.000 | 0.002 | 1.000 | 0.224 | 0.001 | 0.087 |
| 9 | Time mobile | 0.003 | 0.680 | 0.154 | 1.000 | 0.006 | 0.060 | 0.944 |
| 10 | Time immobile | 0.003 | 0.680 | 0.154 | 1.000 | 0.006 | 0.060 | 0.944 |
| 11 | Mobile episodes | 0.000 | 0.259 | 1.000 | 0.792 | 0.002 | 0.032 | 0.274 |
| 12 | Immobile episodes | 0.000 | 0.264 | 1.000 | 0.776 | 0.001 | 0.034 | 1.000 |
| 13 | Mobility latency | 0.747 | 1.000 | 1.000 | 0.142 | 1.000 | 1.000 | 0.305 |
| 14 | Immobility latency | 0.039 | 1.000 | 1.000 | 0.479 | 0.540 | 0.113 | 0.285 |
| 15 | Rotations | 0.027 | 1.000 | 0.296 | 1.000 | 0.006 | 0.758 | 0.623 |
| 16 | Clockwise rotations | 0.533 | 1.000 | 0.016 | 1.000 | 0.037 | 0.236 | 1.000 |
| 17 | Anti-clockwise rotations | 0.363 | 1.000 | 0.107 | 1.000 | 0.999 | 0.765 | 0.157 |
| 18 | Absolute turn angle | 0.003 | 1.000 | 0.276 | 1.000 | 0.064 | 0.029 | 0.141 |
| 19 | Path efficiency | 0.041 | 0.883 | 0.015 | 1.000 | 1.000 | 1.000 | 1.000 |
| 20 | Line crossings | 0.178 | 0.010 | 0.000 | 0.004 | 0.007 | 0.094 | 1.000 |
| **16-grid measure** |  |  |  |  |  |  |  |  |
| 21 | 16-grid crossings | 0.114 | 0.004 | 0.000 | 1.000 | 0.625 | 0.014 | 0.003 |
| **Centre-point measures** |  |  |  |  |  |  |  |  |
| 22 | Mean distance from | 0.000 | 0.085 | 0.037 | 0.105 | 0.002 | 0.789 | 1.000 |
| 23 | Max. distance from | 0.000 | 0.085 | 0.037 | 0.105 | 0.001 | 1.000 | 0.275 |
| 24 | Min. distance from | 0.000 | 0.127 | 0.024 | 1.000 | 0.003 | 0.559 | 0.018 |
| 25 | Time moving towards | 0.149 | 0.000 | 0.001 | 0.000 | 0.618 | 0.265 | 0.655 |
| 26 | Time moving away | 0.001 | 0.037 | 1.000 | 0.000 | 0.023 | 0.149 | 0.704 |
| 27 | Mean speed moving towards | 0.897 | 1.000 | 0.000 | 0.429 | 0.075 | 0.024 | 1.000 |
| 28 | Initial heading error | 0.395 | 1.000 | 1.000 | 1.000 | 1.000 | 0.059 | 0.599 |
| 29 | Average absolute heading error | 0.000 | 0.030 | 0.071 | 0.325 | 1.000 | 1.000 | 1.000 |
| **Thigmotaxis measures** |  |  |  |  |  |  |  |  |
| 30 | Entries | 0.174 | 0.010 | 0.000 | 0.005 | 0.007 | 0.093 | 1.000 |
| 31 | Number exits | 0.179 | 0.010 | 0.000 | 0.004 | 0.007 | 0.094 | 0.550 |
| 32 | Time | 0.000 | 0.055 | 0.042 | 0.011 | 0.002 | 0.435 | 0.442 |
| 33 | Was 1st zone | 1.000 | 1.000 | 0.698 | 1.000 | 1.000 | 0.151 | 1.000 |
| 34 | Distance | 0.000 | 0.000 | 0.000 | 0.000 | 0.448 | 0.279 | 0.067 |
| 35 | Distance to first entry | 0.166 | 1.000 | 0.019 | 0.787 | 0.141 | 0.611 | 0.218 |
| 36 | Latency to first entry | 0.232 | 1.000 | 0.012 | 1.000 | 0.013 | 0.641 | 0.063 |
| 37 | Latency to first exit | 0.334 | 1.000 | 0.016 | 1.000 | 0.565 | 0.618 | 0.026 |
| 38 | Latency to last entry | 0.211 | 0.478 | 0.567 | 1.000 | 0.656 | 0.443 | 0.182 |
| 39 | Average speed | 0.001 | 0.000 | 0.000 | 0.007 | 0.163 | 0.063 | 0.094 |
| 40 | Max. speed | 0.000 | 0.860 | 0.000 | 1.000 | 0.020 | 0.729 | 0.018 |
| 41 | Max. visit | 1.000 | 0.499 | 0.184 | 0.108 | 1.000 | 0.541 | 0.444 |
| 42 | Min. visit | 1.000 | 0.721 | 1.000 | 1.000 | 1.000 | 1.000 | 1.000 |
| 43 | Mean visit | 0.922 | 0.003 | 1.000 | 0.102 | 0.009 | 0.124 | 1.000 |
| 44 | Time mobile | 0.000 | 0.002 | 0.000 | 0.001 | 0.218 | 0.756 | 0.318 |
| 45 | Time immobile | 0.008 | 1.000 | 0.856 | 0.400 | 0.004 | 0.169 | 0.984 |
| 46 | Immobile episodes | 0.000 | 0.839 | 0.069 | 0.874 | 0.015 | 0.081 | 1.000 |
| 47 | Initial distance | 0.462 | 1.000 | 1.000 | 0.260 | 0.033 | 0.099 | 0.002 |
| 48 | Mean distance from | 0.000 | 1.000 | 0.031 | 1.000 | 0.001 | 1.000 | 1.000 |
| 49 | Max. distance from | 1.000 | 0.055 | 1.000 | 0.249 | 1.000 | 1.000 | 0.147 |
| 50 | Min. distance from | 1.000 | 1.000 | 0.698 | 1.000 | 1.000 | 0.151 | 1.000 |
| 51 | Cumulative distance | 0.000 | 0.339 | 0.042 | 0.497 | 0.001 | 0.833 | 0.326 |
| 52 | Mean distance to border | 0.023 | 0.319 | 1.000 | 0.432 | 1.000 | 1.000 | 1.000 |
| 53 | Max. distance to border | 0.029 | 1.000 | 0.349 | 1.000 | 1.000 | 1.000 | 1.000 |
| 54 | Min. distance to border | 1.000 | 1.000 | 1.000 | 1.000 | 1.000 | 1.000 | 1.000 |
| 55 | Time getting closer to zone | 0.232 | 1.000 | 0.230 | 1.000 | 0.014 | 0.414 | 0.043 |
| 56 | Time getting further from zone | 0.253 | 1.000 | 0.248 | 1.000 | 0.057 | 0.423 | 0.101 |
| 57 | Initial heading error | 0.277 | 0.953 | 1.000 | 1.000 | 0.868 | 0.144 | 0.416 |
| 58 | Signed initial heading error | 1.000 | 1.000 | 1.000 | 1.000 | 0.563 | 0.711 | 0.514 |
| 59 | Average absolute heading error | 0.001 | 0.001 | 0.000 | 0.000 | 1.000 | 1.000 | 1.000 |
| 60 | Time moving towards | 0.021 | 0.000 | 0.893 | 0.000 | 0.850 | 0.350 | 0.636 |
| 61 | Time moving away from | 0.000 | 0.844 | 0.093 | 0.016 | 0.011 | 0.253 | 0.497 |
| 62 | In zone oriented to centre | 0.005 | 0.005 | 0.354 | 0.467 | 0.867 | 0.480 | 0.158 |
| 63 | Absolute turn angle | 0.000 | 0.009 | 0.000 | 0.073 | 0.226 | 0.208 | 0.040 |
| 64 | Freezing episodes | 0.049 | 0.131 | 0.562 | 1.000 | 0.061 | 1.000 | 0.399 |
| 65 | Time freezing | 0.131 | 0.296 | 1.000 | 0.515 | 0.066 | 0.612 | 0.532 |
| 66 | Path efficiency to entry | 0.523 | 0.686 | 0.241 | 1.000 | 0.991 | 0.746 | 0.480 |
| 67 | Corrected integrated path length | 0.000 | 0.336 | 0.040 | 0.464 | 0.001 | 0.858 | 0.348 |
| 68 | Number of line crossings | 0.177 | 0.010 | 0.000 | 0.005 | 0.007 | 0.093 | 1.000 |

<sup>a</sup>Analysis of variance with post-hoc Bonferroni's corrections. <sup>b</sup>Independent T-test analysis, associated heat-map representations in Figure 2. *Experiment 1*, with no head-stage/NAT-1; *Experiment 2*, with head-stage/NAT-1; ref., referenced to; Max., maximum; Min., minimum.

**Table S2.** Statistical summary of comparisons between treatment groups in Experiments 1 and 2 in pre-selected parameters. Conventional statistical comparisons against saline which differ between experimental groups are highlighted in red- significance is considered where p<0.05. All statistical analysis for comparisons to saline in each experimental group were resampled 1000 (seed starting at 123456) in R (v.4.3.0).

| Parameters |  | Within-experiment comparisons<sup>a</sup> |  |  |  |  |  | MK801 minus saline |  | Scopolamine minus saline |  |
| --- | --- | --- | --- | --- | --- | --- | --- | --- | --- | --- | --- |
| | | Saline | MK801 | Scopolamine | F<sup>b</sup> | Sig. | $\eta p^2$ | Mean difference<br>[95% CI] | $p^a$ | Mean difference<br>[95% CI] | $p^a$ |
|  |  | mean±SD | mean±SD | mean±SD |  |  |  |  |  |  |  |
| Distance moved (m) | E1 | 131.6±20.2 | 70.0±25.6 | 182.2±20.0 | 61.9 | <0.001 | 0.826 | -61.6 [-81.5; -42.2] | <0.001 | 50.6 [34.1; 68] | <0.001 |
|  | E2 | 139.9±19.2 | 39.3±23.5 | 155.2±26.7 | 65.3 | <0.001 | 0.845 | -101 [-120; -79.7] | <0.001 | 15.3 [-8.76; 32.5] | 0.531 |
| Average distance from centre-point (m) | E1 | 0.218±0.007 | 0.155±0.03 | 0.237±0.013 | 48.9 | <0.001 | 0.790 | -0.065 [-0.080; -0.044] | <0.001 | 0.020 [0.008; 0.026] | 0.085 |
|  | E2 | 0.199±0.01 | 0.161±0.050 | 0.230±0.012 | 12.2 | <0.001 | 0.505 | -0.039 [-0.082; -0.014] | 0.037 | 0.032 [0.019; 0.042] | 0.105 |
| Thigmotaxis (ratio) | E1 | 0.532±0.052 | 0.223±0.157 | 0.807±0.147 | 50.5 | <0.001 | 0.795 | -0.308 [-0.403; -0.201] | <0.001 | 0.278 [0.118; 0.339] | <0.001 |
|  | E2 | 0.463±0.049 | 0.285±0.189 | 0.777±0.128 | 30.7 | <0.001 | 0.710 | -0.179 [-0.304; -0.062] | 0.038 | 0.316 [0.219; 0.394] | <0.001 |
| Thigmotaxis (s) | E1 | 1266.1±96.3 | 545.1±379.3 | 1539.8±184.5 | 41.7 | <0.001 | 0.762 | -721 [-927; -473] | <0.001 | 274 [95.9; 363] | 0.055 |
|  | E2 | 1050.0±105.7 | 699.2±445.6 | 1475.8±163.8 | 17.3 | <0.001 | 0.590 | -351 [-623; -75.8] | 0.042 | 426 [280; 538] | 0.011 |
| Line-crossings | E1 | 1060.6±145.3 | 866.9±265.1 | 1374.2±154.3 | 16.9 | <0.001 | 0.565 | -194 [-403; -19.2] | 0.114 | 314 [187; 440] | 0.004 |
|  | E2 | 1030.0±132.49 | 458.3±238.5 | 1012.0±196.9 | 25.4 | <0.001 | 0.679 | -574 [-724; -371] | <0.001 | -20 [-196; 97.6] | >0.999 |
| Meandering (degrees/m) | E1 | 912.5±89.5 | 2716.4±1008.4 | 705.6±73.5 | 35.9 | <0.001 | 0.734 | 1800 [1260; 2480] | <0.001 | -207 [-277; -142] | >0.999 |
|  | E2 | 932.0±107.4 | 3077±1813.4 | 750.4±89.2 | 13.7 | <0.001 | 0.533 | 2150 [1330; 3860] | 0.001 | -182 [-264; -98.3] | >0.999 |
| Rotation | E1 | 76.5±10.5 | 180.1±142.7 | 99.8±10.6 | 4.4 | 0.023 | 0.251 | 104 [27.4; 209] | 0.027 | 23.3 [14.7; 32.1] | >0.999 |
|  | E2 | 98.8±12.5 | 158.2±138.3 | 96.1±18.2 | 1.9 | 0.176 | 0.135 | 65.4 [-0.778; 175] | 0.293 | 3.33 [-11.3; 15.6] | >0.999 |

<sup>a</sup>Bootstrapped analysis of variance with Bonferroni's corrections. <sup>b</sup>Experiment 1, F(2,26); Experiment 2, F(2,24). *SD*, standard deviation; *F*, *f*-values; sig., *p*-values with significance levels set at $p < 0.05$ ; $\eta p^2$ , partial eta-squared; CI, confidence intervals (lower; upper bounds); E1, Experiment 1 with no head-stage/NAT-1; and E2, Experiment 2 with head-stage/NAT-1.

